# Tumor Suppression across the Human miRNome Associates with Guanine-Rich Precursor Terminal Loops

**DOI:** 10.64898/2026.01.26.701701

**Authors:** Amit Cohen, Mario Alberto Burgos-Aceves, Yoav Smith

## Abstract

MicroRNAs (miRNAs) play essential regulatory roles in controlling cell growth, proliferation, and differentiation in cancer. While functional studies have identified numerous oncogenic (oncomiRs) and tumor suppressor (TS) miRNAs, the structural features that differentiate these groups remain poorly understood. Here, we performed a comprehensive sequence analysis of 955 human pre-miRNA terminal loops (TLs), focusing on enrichment of single guanine (G) and GG dinucleotides. A quantitative G enrichment score was used to define 42 G-rich TL miRNAs and 17 G-free TL miRNAs as controls. Functional roles of these miRNAs were curated from 757 publications. The results show that G-rich TL miRNAs consistently display higher TS/oncomiR ratios than G-free TL miRNAs across most cancer types, with a significant enrichment observed in lung cancer (p = 0.023). Focusing on miR-139, a TS miRNA with a G-rich TL, integrative analysis of publicly available transcriptomic and proteomic data revealed its consistent downregulation across all stages of lung adenocarcinoma (LUAD), accompanied by reciprocal overexpression of its validated oncogenic target, CCNB1. These findings highlight the biological relevance of G-rich TL structures in miRNA-mediated tumor suppression in lung cancer, suggesting their consideration in future therapeutic strategies aimed at restoring vulnerable TS miRNAs.

## Introduction

MicroRNAs (miRNAs) are short non-coding RNAs that post-transcriptionally regulate gene expression by guiding the RNA-induced silencing complex (RISC) to complementary mRNA targets [1]. Through their ability to modulate multiple transcripts simultaneously, miRNAs function as key regulators of oncogenic and tumor-suppressive regulatory networks across human cancers [2]. Although extensive functional studies have classified numerous miRNAs as oncogenic (oncomiRs) or tumor suppressor (TS), the structural determinants that predispose a miRNA toward either activity remain incompletely defined.

One structural feature of growing interest is the terminal loop (TL) of the precursor miRNA (pre-miRNA). Rather than serving merely as a connector between the hairpin arms, the TL is a dynamic hub involved in miRNA biogenesis [3]. TL sequence and structure influence recognition by the Microprocessor complex (Drosha–DGCR8), modulate Dicer cleavage precision, and mediate interactions with RNA-binding proteins (RBPs) that selectively enhance or inhibit miRNA maturation [4]. Even subtle changes in loop composition can markedly alter pre-miRNA folding, accessibility, and processing efficiency [5, 6].

Among TL sequence features, guanine (G) enrichment is particularly noteworthy. G has the lowest oxidation potential of the four RNA bases, and the most deleterious oxidized base in RNA is 8-hydroxyguanosine (8-oxoG) [7, 8]. Such modifications preferentially occur in exposed RNA regions [9], including TLs, as observed in the yeast 5S rRNA [10, 11]. These biochemical properties suggest that G-rich TLs may be functionally sensitive to redox conditions and particularly vulnerable in tissues with high oxidative stress.

The lung is one of the most oxidatively stressed organs in the human body. Chronic exposure to cigarette smoke, airborne pollutants, inflammatory infiltrates, and metabolic ROS collectively generates a redox environment far more extreme than that of most tissues [12]. This environment also contributes to the well-documented global downregulation of miRNAs in smokers’ lungs [13–16]. Thus, the lung represents a biologically plausible setting in which structurally vulnerable miRNAs, such as those with G-rich TLs, might undergo preferential degradation or impaired processing.

Previous studies from our group provided initial evidence that TL G content may have biological significance, as G enrichment in pre-miRNA TLs correlated with broad miRNA suppression in cancer [17], and G-rich TLs exhibited a trend toward TS activity [18]. However, these analyses focused on selected miRNA subsets and did not address whether G-rich TLs constitute a functionally coherent class across the human miRNome.

In the present work, we comprehensively analyze all 955 human pre-miRNA TLs, quantify G enrichment using an empirical distribution-based scoring system, and systematically integrate functional annotation from 757 curated publications. Our results reveal that in lung cancer, TS miRNAs exhibit significantly higher G-richness in their TLs compared to oncomiRs, and that miR-139, one of the most strongly TS miRNAs with a G-rich TL, is consistently downregulated in lung adenocarcinoma (LUAD), pointing to a potential mechanistic link between TL composition and TS miRNA loss in this highly oxidative tissue environment.

## Materials and Methods

### Dataset Collection

All human pre-miRNA, 5’ mature, and 3’ mature sequences were retrieved from the Sanger Institute miRBase database version 22 (http://microrna.sanger.ac.uk/sequences/). TL sequences were extracted by omitting the 5’ and 3’ mature sequences from the pre-miRNA sequence.

### Quantification of Guanine Content in TLs

Nucleotide composition, including G single-nucleotide and GG dinucleotide frequencies, in miRNA TLs was calculated using the compseq algorithm (http://emboss.bioinformatics.nl/cgi-bin/emboss/compseq).

### Literature Data Mining

Literature searches were performed in the PubMed literature database for original articles written in the English language, focusing on miRNAs and cancer. The searches included a specific miRNA term paired with the keyword ‘cancer’. Restriction was set for the publication date ranging from 2015 to 2025. Only articles showing a role for miRNAs, by using functional studies, were selected. For a total of 42 G-rich TL miRNAs and 17 G-free TL miRNAs, we manually curated 757 publications. Where applicable, the cancer type and experimentally validated target genes were retrieved from the above articles.

### Pathway Analysis of G-Rich TL miRNA Targets in Lung Cancer

Experimentally validated targets of G-rich TL miRNAs, retrieved from curated lung cancer publications, were analyzed for KEGG pathway enrichment using the g:Profiler tool (https://biit.cs.ut.ee/gprofiler/gost), and their known protein–protein interactions were visualized using STRING v12 (http://string-db.org/).

### Genomic Data Analysis

Genomic computational analysis involved the investigation of open-source datasets to examine the expression profiles of hsa-miR-139 in both healthy and cancerous lung tissues. For this analysis, data were utilized from The Cancer Genome Atlas (TCGA) (<u>The Cancer Genome Atlas Program (TCGA) - NCI</u>). CCNB1 protein expression data were retrieved from the Clinical Proteomic Tumor Analysis Consortium (CPTAC) database (<u>Clinical Proteomic Tumor Analysis Consortium (CPTAC) | NCI Genomic DataCommons</u>). Normalized CCNB1 protein abundance values were analyzed across all available CPTAC samples, including normal lung tissue and primary tumors stratified by pathological stage. Prior to visualization, values were standardized across the entire cohort using Z-score normalization to enable comparison of relative expression levels.

### G Enrichment Scoring

To score G enrichment we calculated a composite metric that integrates single-G and double-G frequencies within the precursor’s TL according to the following formula:

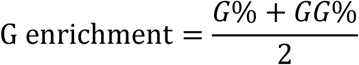

where G% and GG% denote the relative frequencies of single and consecutive double G bases, respectively. These metrics were used to calculate a G enrichment score across the entire miRNome.

### Empirical p-value Estimation

Empirical p-values were calculated for each miRNA to assess the statistical significance of TL G enrichment. For a given miRNA i with a G enrichment score GEᵢ, its p-value was defined as:
pᵢ = (number of miRNAs with G enrichment ≥ GEᵢ) / N,
where N = 955 (the total number of pre-miRNAs analyzed).

### Statistical Data Analysis

To assess whether the effect of G enrichment on TS/oncomiR ratios differed between tissues, we tested for interaction between G-status (G-rich vs. G-free) and tissue type using the Breslow–Day test for homogeneity of odds ratios. This test evaluates whether the odds ratio of TS versus oncomiRs for G-rich compared with G-free TL miRNAs differs significantly across cancer types relative to the “All Cancers” background. A p-value < 0.05 was considered statistically significant, and all tests were two-tailed.

A two-tailed t-test was performed on the gene expression data for lung cancer vs. normal lungs. Data distributions were visualized using standard box-and-whisker plots, in which the central line represents the median, the box corresponds to the interquartile range (IQR), and the whiskers extend to 1.5× IQR.

## Results

Since G enrichment in pre-miRNA TLs was previously associated with global miRNA downregulation and TS function in cancer [17, 18], we asked whether this structural feature is consistently linked to the functional classification of miRNAs across the entire human miRNome. For this purpose, human miRNA sequences (pre-miRNA, 5’ mature, and 3’ mature) were retrieved from the miRBase database. TL sequences were extracted by removing the 5’ and 3’ mature miRNA regions from the full pre-miRNA sequences, yielding a total of 955 TL segments. To systematically evaluate G enrichment, we first quantified G and GG content across all human precursor miRNA TL sequences (Supplementary Table 1). In addition, we applied a standardized scoring metric to identify G enrichment within the TLs; a composite G enrichment score was calculated for each pre-miRNA TL by averaging the frequencies of single G bases (G%) and consecutive G pairs (GG%) within the TL, and an empirical p-value reflecting TL G enrichment was assigned to each miRNA (Supplementary Table 1). This strategy allowed us to define a G-rich set and assess whether loop G composition aligns with known functional activity. Based on their empirical G enrichment values, 42 miRNAs were assigned as the group with the most G-rich TLs (Table 1), and 17 miRNAs representing the G-free TLs served as a control group (Supplementary Table 2).

**Table 1.** A list of 42 miRNAs with a significant pre-miRNA G-rich TL sequences. Presented are G and GG percents and G enrichment empirical p-values.

| **miRNA** | **G%** | **GG%** | **Empirical p-value** |
| --- | --- | --- | --- |
| hsa-miR-10400 | 0.67 | 0.45 | 0.001 |
| hsa-miR-642a | 0.62 | 0.50 | 0.002 |
| hsa-miR-6850 | 0.67 | 0.43 | 0.003 |
| hsa-miR-4717 | 0.60 | 0.44 | 0.004 |
| hsa-miR-6819 | 0.62 | 0.42 | 0.005 |
| hsa-miR-320a | 0.60 | 0.33 | 0.006 |
| hsa-miR-6871 | 0.57 | 0.33 | 0.007 |
| hsa-miR-6790 | 0.53 | 0.36 | 0.008 |
| hsa-miR-132 | 0.57 | 0.31 | 0.009 |
| hsa-miR-3928 | 0.55 | 0.30 | 0.011 |
| hsa-miR-939 | 0.55 | 0.30 | 0.011 |
| hsa-miR-6825 | 0.50 | 0.33 | 0.013 |
| hsa-miR-6722 | 0.56 | 0.27 | 0.015 |
| hsa-miR-1233 | 0.51 | 0.31 | 0.016 |
| hsa-miR-6890 | 0.53 | 0.29 | 0.018 |
| hsa-miR-3619 | 0.56 | 0.25 | 0.019 |
| hsa-miR-345 | 0.57 | 0.23 | 0.020 |
| hsa-miR-5088 | 0.46 | 0.33 | 0.021 |
| hsa-miR-3691 | 0.50 | 0.29 | 0.022 |
| hsa-miR-9851 | 0.54 | 0.25 | 0.024 |
| hsa-miR-139 | 0.54 | 0.25 | 0.024 |
| hsa-miR-6720 | 0.54 | 0.25 | 0.024 |
| hsa-miR-1229 | 0.45 | 0.33 | 0.026 |
| hsa-miR-6765 | 0.54 | 0.25 | 0.027 |
| hsa-miR-1228 | 0.53 | 0.24 | 0.028 |
| hsa-miR-193a | 0.50 | 0.27 | 0.029 |
| hsa-miR-6847 | 0.50 | 0.27 | 0.030 |
| hsa-miR-6743 | 0.50 | 0.26 | 0.031 |
| hsa-miR-6808 | 0.45 | 0.30 | 0.033 |
| hsa-miR-4746 | 0.45 | 0.30 | 0.033 |
| hsa-miR-1207 | 0.48 | 0.27 | 0.034 |
| hsa-miR-4787 | 0.53 | 0.21 | 0.035 |
| hsa-miR-3677 | 0.43 | 0.31 | 0.036 |
| hsa-miR-6800 | 0.45 | 0.28 | 0.037 |
| hsa-miR-6511b | 0.50 | 0.24 | 0.041 |
| hsa-miR-6511a | 0.50 | 0.24 | 0.041 |
| hsa-miR-4747 | 0.40 | 0.33 | 0.045 |
| hsa-miR-33b | 0.56 | 0.18 | 0.046 |
| hsa-miR-4757 | 0.50 | 0.23 | 0.047 |
| hsa-miR-1301 | 0.50 | 0.23 | 0.047 |
| hsa-miR-4758 | 0.48 | 0.25 | 0.049 |
| hsa-miR-4708 | 0.50 | 0.22 | 0.050 |

To assess whether G enrichment correlates with known biological function, we systematically mined the literature for studies published in PubMed database between 2015 and 2025 that functionally characterized the TS or oncomiR roles of these miRNAs. A total of 757 publications were curated (Supplementary Table 2). Cancer type information was also retrieved from these publications.

Across the “All Cancers” dataset (757 publications), G enrichment in pre-miRNA TLs was associated with a higher TS/oncomiR ratio compared with G-free controls (Figure 1). To determine whether this effect varied across tissues, we tested for interaction between G-status (G-rich vs. G-free) and cancer type using the Breslow–Day test. A significant deviation from homogeneity was detected only in lung cancer (p = 0.023), indicating that the increase in TS/O among G-rich miRNAs was substantially stronger in lung cancer relative to the “All Cancers” background (Figure 1). Head & neck cancer (combining oral, tongue, nasopharyngeal and laryngeal cancers) showed a borderline trend (p = 0.071) but did not reach statistical significance. No significant deviation from the background pattern was observed in any of the other cancer types examined, including breast, cervical, colorectal, esophageal, gastric, glioma, hepatic, ovarian, pancreatic, and prostate cancers.

**Figure 1:**
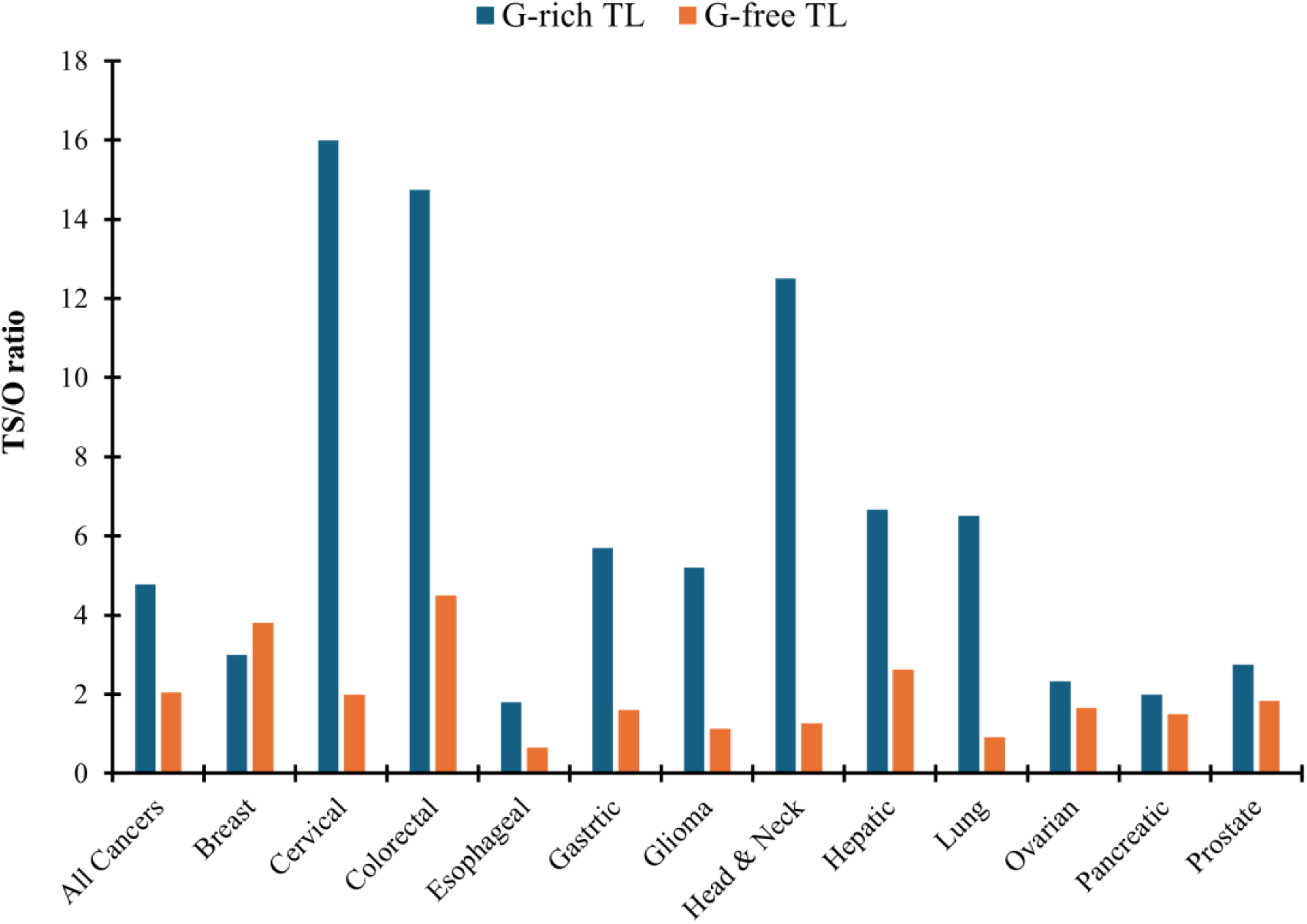
TS/oncomiR ratios of G-rich versus G-free TL miRNAs across human cancers. The TS/oncomiR ratio (TS/O ratio) was calculated based on the number of PubMed-indexed publications (2015–2025) reporting experimentally validated TS versus oncomiR functions for each miRNA. The “All Cancers” category represents the combined dataset of all analyzed cancer types, comprising a total of 757 publications.

G-rich TL miRNAs in lung cancer regulate a functionally coherent network of target genes. In total, we retrieved 61 unique target genes from 75 publications related to the G-rich TL miRNA group and lung cancer, including non-small cell lung cancer (NSCLC) and LUAD (Figure 2A; Supplementary Table 3). KEGG pathway analysis using g:Profiler tool reveals that the miRNA target genes are broadly involved in Proteoglycans in cancer, HIF-1 and ErbB signaling, and endocrine resistance pathways (Figure 2B). Several of these genes serve as common targets in lung cancer for multiple miRNAs. Examples are the miR-139 target genes IGF1R and MMP2, which are also verified targets of miR-320a and miR-1207, respectively (Supplementary Table 3).

**Figure 2:**
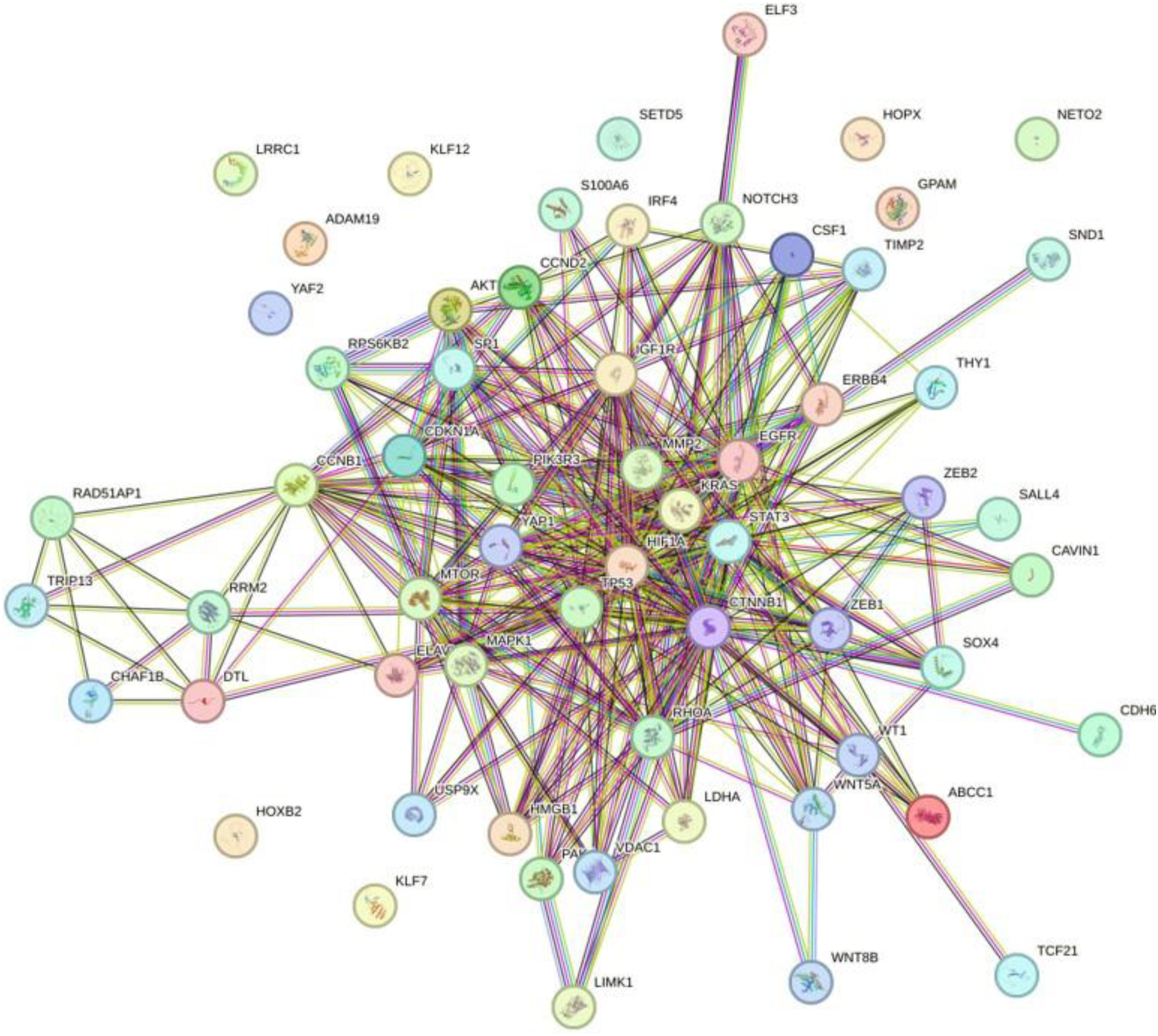
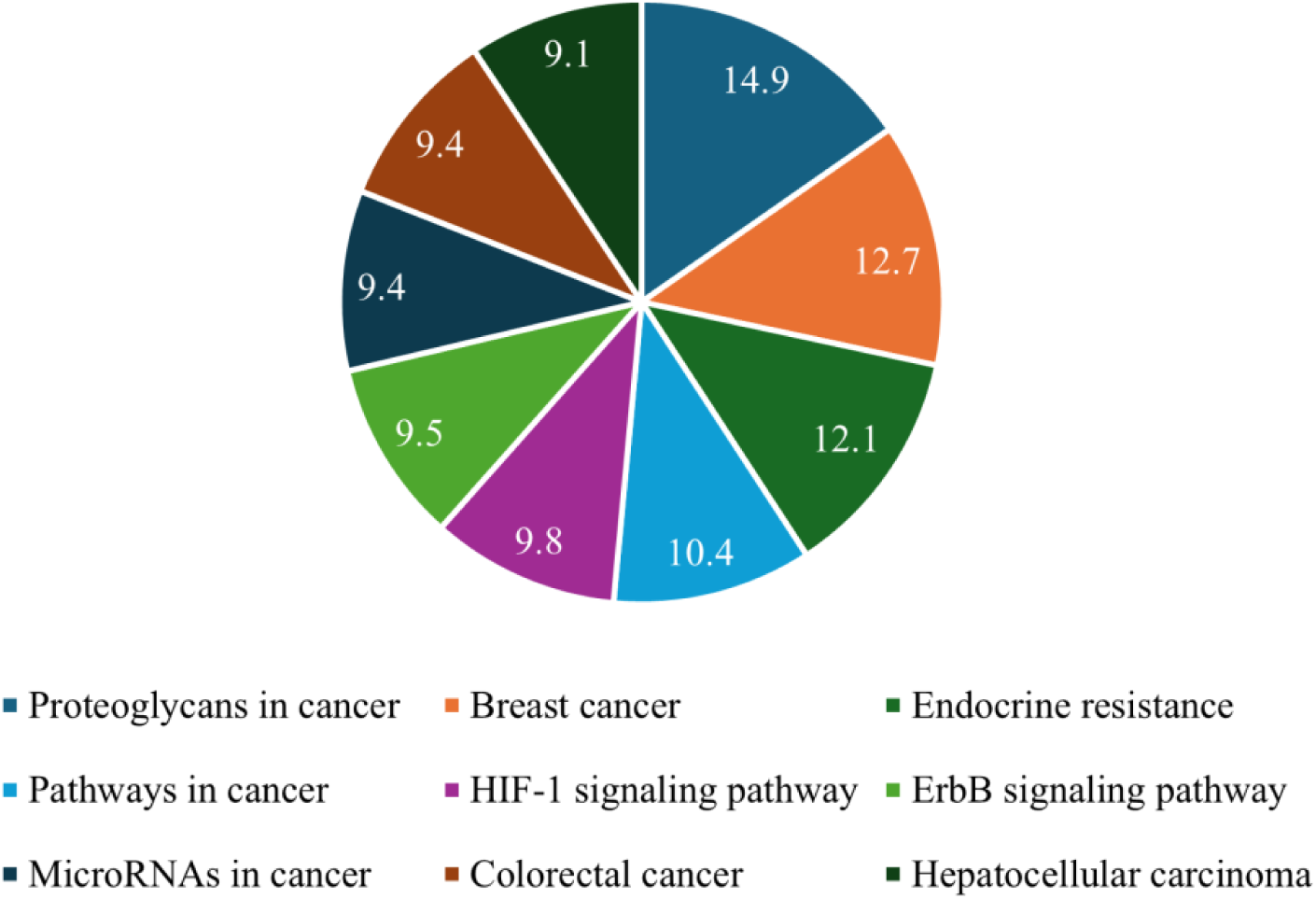
Pathway analysis of G-rich TL miRNAs target genes in lung cancer. **A.** Network connections between the 61 G-rich TL miRNAs target genes, as identified and visualized by the STRING database. **B.** Enriched KEGG pathway terms of 61 target genes of G-rich TL miRNAs in lung cancer. Shown are significant –log2 adjusted p-values, as were identified using the g:Profiler tool.

Since all 42 publications referring to miR-139 consistently describe it as having a TS function, and its pre-miRNA TL sequence (GUGGCUCGGAGGC) is significantly G-rich (54% G and 25% GG; p = 0.024) and contains a conserved RBP-recognition motif (GGAG), we further analyzed its expression across different lung cancer TCGA datasets [19]. Large-scale cancer sample analysis (n=447) shows that miR-139 expression is significantly reduced in primary tumors of LUAD relative to normal tissue (Figure 3A; p = 0.0029). Further analysis of this LUAD sample cohort according to individual cancer stage reveals that miR-139 expression is significantly reduced in all four stages (Stage 1, 2, 3 and 4) relative to normal tissue (Figure 3B; p = 0.0029, p = 0.0027, p = 0.0026 and p = 0.009, respectively).

**Figure 3:**
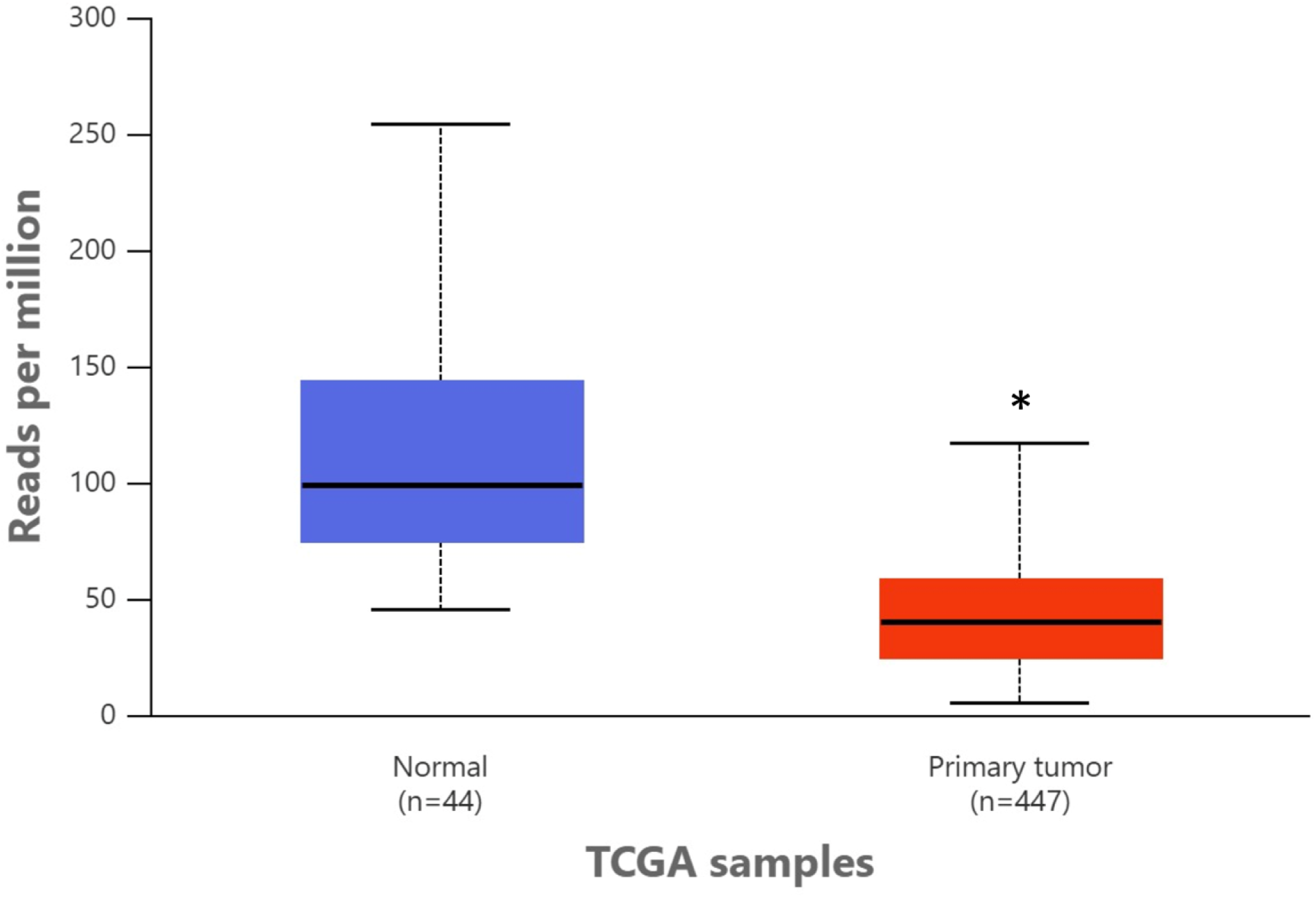
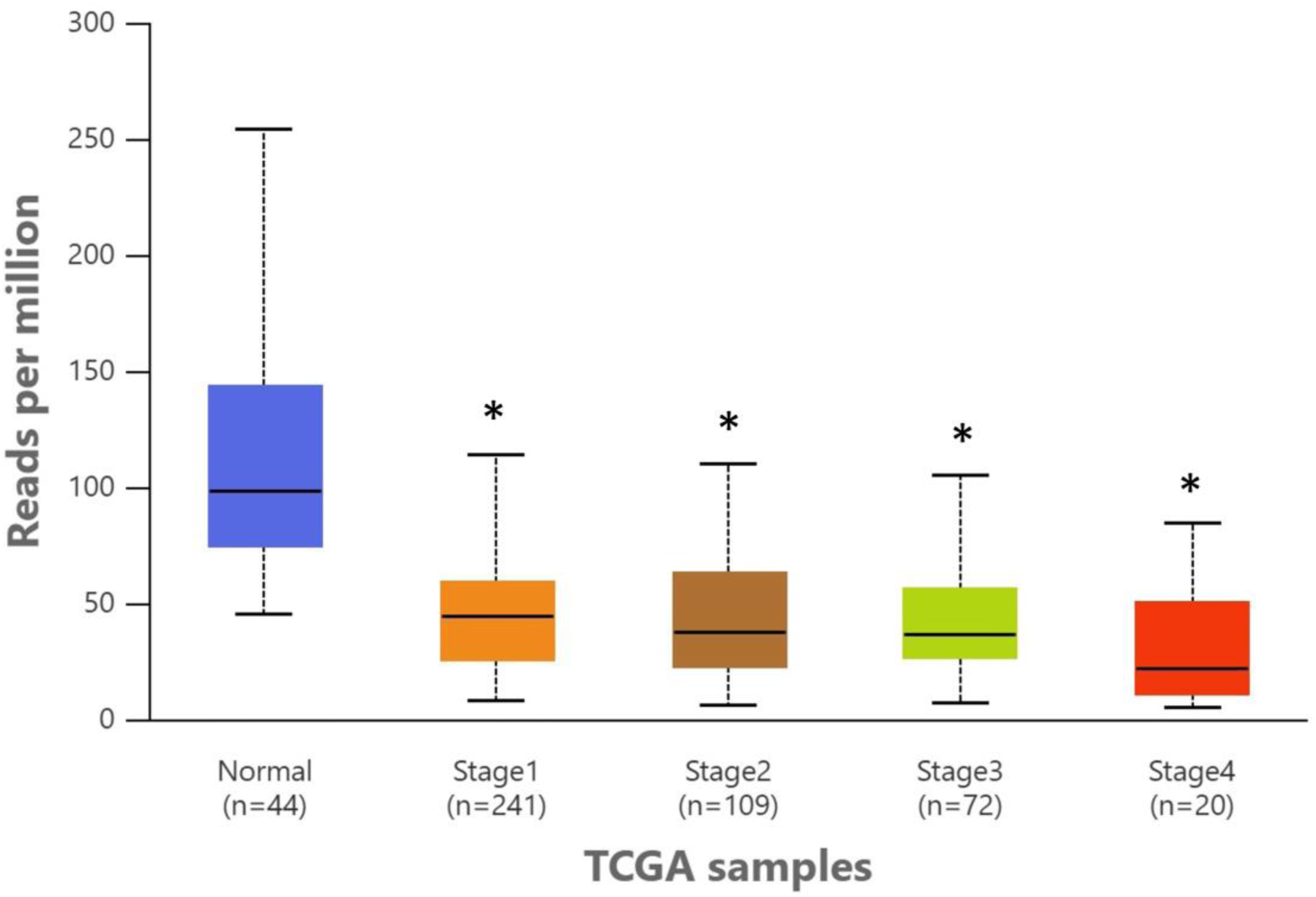
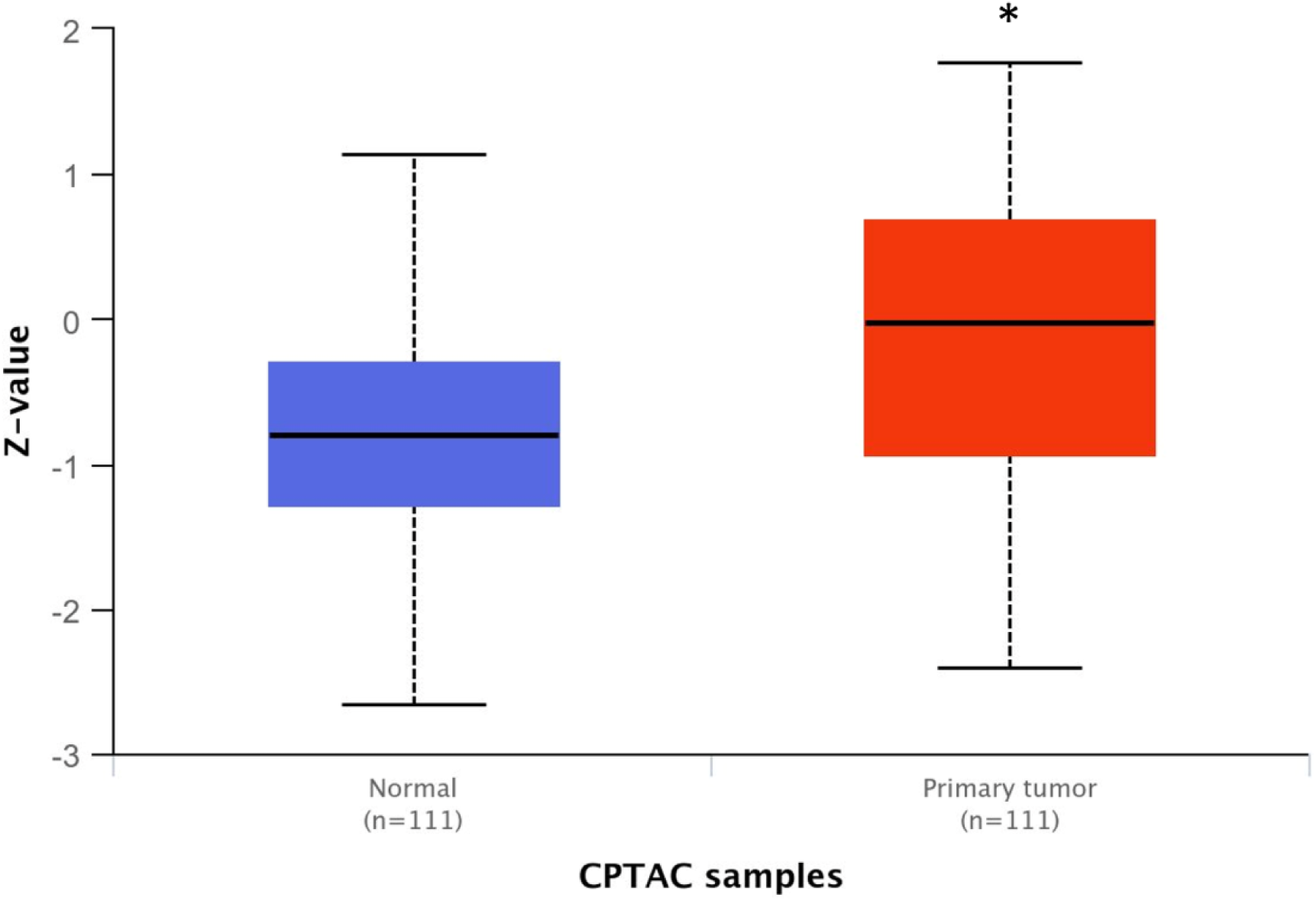
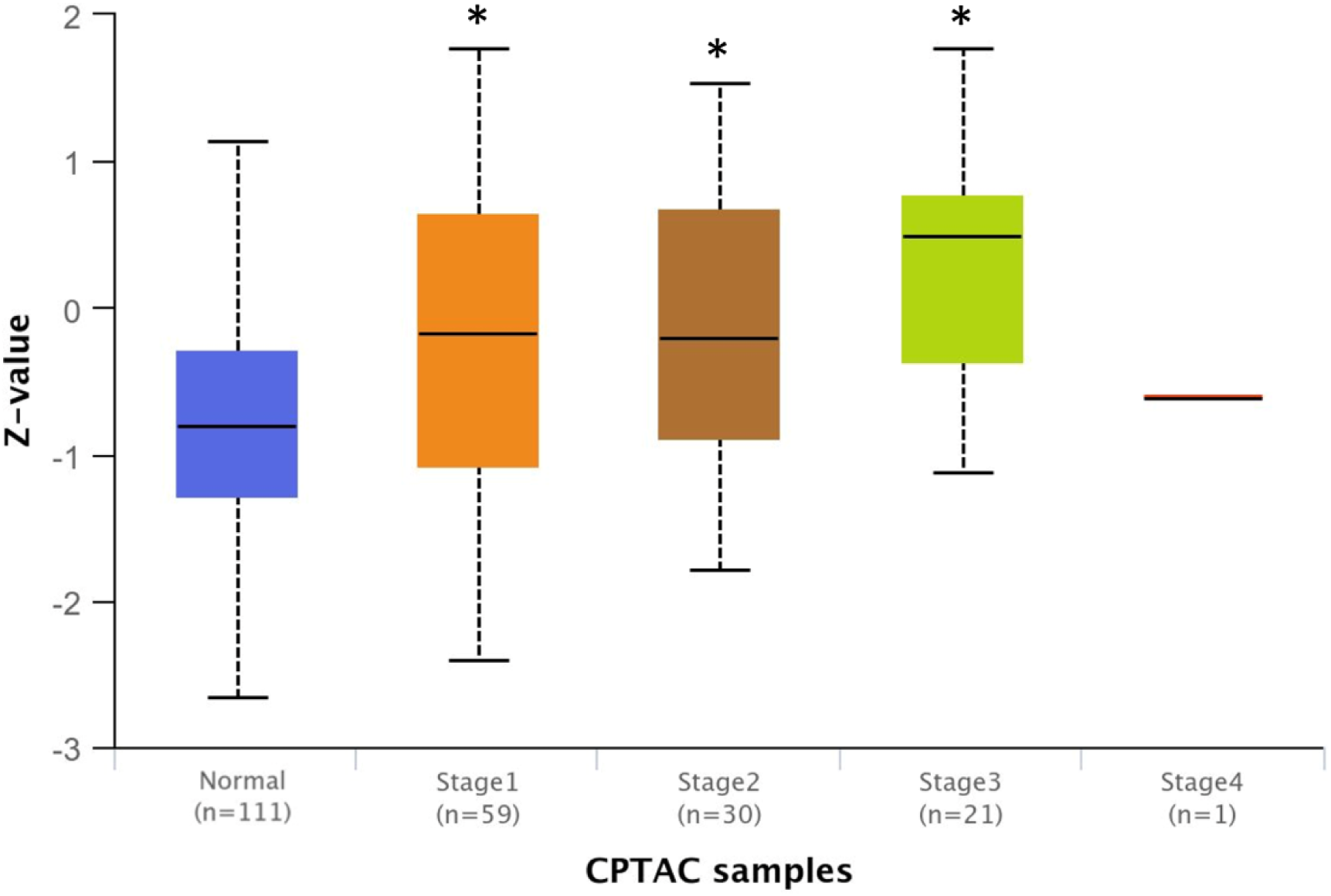
Expression of miR-139 and CCNB1 in LUAD. **A.** miR-139 expression levels in normal and LUAD primary tumors. **B.** miR-139 expression levels in normal and different cancer stages (stage 1, 2, 3, and 4) of LUAD primary tumors. Analysis was conducted on data retrieved from the TCGA database. Sample numbers are indicated beneath each column. *p<0.05. **C.** CCNB1 protein expression levels in normal and LUAD primary tumors. **D.** CCNB1 protein expression levels in normal and different cancer stages (stage 1, 2, 3, and 4) of LUAD primary tumors. Analysis was conducted on data retrieved from the CPTAC database. Sample numbers are indicated beneath each column. *p<0.05. The y-axis represents Z-score–normalized CCNB1 protein expression values calculated across samples.

One of the confirmed target genes of miR-139 in LUAD is Cyclin B1 (CCNB1) [20]. To investigate further, CCNB1 protein expression data in LUAD (n=111) were obtained from the CPTAC database and analyzed similarly to the miR-139 expression data. The results show an inverse pattern compared to those of miR-139: CCNB1 protein expression is significantly elevated in primary tumors of LUAD compared to normal tissue (Figure 3C; p = 1.2×10^−6^). Further analysis of this LUAD sample cohort by individual cancer stage shows that CCNB1 expression is significantly increased across three stages (Stage 1, 2 and 3) relative to normal tissue (Figure 3D; p = 0.0017, p = 0.0015 and p = 3.3×10^−5^, respectively). CCNB1 expression in stage 3 was also significantly higher than in stage 1 (Figure 3D; p = 0.041).

## DISCUSSION

Global repression of miRNA expression is a common molecular characteristic of cancer [21]. While previous work from our group showed that G-rich TLs are disproportionately found among downregulated miRNAs in cancer and enriched in TS miRNAs [17, 18], the current analysis extends these findings: it creates and analyzes an entire set of 955 human miRNA TLs, establishes an empirical, distribution-based G enrichment score, and systematically integrates functional evidence from a comprehensive set of experimentally validated publications.

A notable example emerging from our analysis is miR-139, which displays one of the highest TL G enrichment scores and is consistently characterized as a TS across 42 independent publications. Consistent with its structural classification, we demonstrate that miR-139 is significantly downregulated in LUAD across all stages of cancer. Its experimentally confirmed target, CCNB1, a key regulator of the G2/M transition and a well-established oncogenic driver [22], displays the opposite pattern: it is strongly upregulated across LUAD tumors and increases with disease stage. This inverse relationship reinforces the functional importance of the miR-139/CCNB1 axis in lung tumorigenesis. Because CCNB1 overexpression is repeatedly linked to poor patient outcomes [23, 24], reduced expression of miR-139 provides a suggested biological route by which TS G-rich TL miRNAs contribute to suppression of cancer growth under normal conditions but are lost in lung cancer.

Target-gene enrichment analysis further shows that G-rich TL miRNAs regulate in lung cancer a tightly interconnected oncogenic network involving KRAS, IGF1R, ERBB4, MMP2, SOX4, KLF7, and others - many of which were previously identified as canonical drivers of lung tumorigenesis, therapy resistance, and metastasis [25–30]. The recurrence of certain targets across multiple independent miRNAs (e.g., IGF1R targeted by both miR-139 and miR-320a) suggests that G-rich TL miRNAs function to buffer against oncogenic activation in coordinated TS modules, similar to the synergistic function previously described in lung cancer for miR-34a and miR-15a/16 cluster [31]. The heightened susceptibility of G-rich TLs to oxidative disruption could therefore explain why multiple TS miRNAs are simultaneously lost in lung cancer, enabling deregulation of the same oncogenic pathways through convergent mechanisms. The association between G-rich TL of miRNA and TS activity in lung cancer aligns with the unique oxidative environment of lung tissue: it is chronically exposed to exogenous and endogenous ROS from cigarette smoke, particulate air pollution, chronic inflammation, and mitochondrial dysfunction [32, 33]. Because G has the lowest oxidation potential of all nucleobases, G-rich-exposed RNA loops are particularly susceptible to oxidative lesions such as 8-oxoG, a modification that can potentially impair miRNA processing [34, 35].

These biochemical vulnerabilities provide a mechanistic explanation for the preferential loss of TS miRNAs and the induction of their oncogenic target genes. A possible model for these phenomena is that the striking susceptibility of G to oxidation in pre-miRNA TLs may reflect not merely biochemical vulnerability but potentially a regulatory design that leverages cellular redox fluctuations. Because of the lowest oxidation potential of G, its enrichment in the exposed TL could allow specific pre-miRNAs to act as redox-responsive switches whose processing efficiency shifts with intracellular ROS levels. Under normal physiological conditions, transient oxidative bursts may temporarily modulate the maturation of G-rich TL miRNAs, enabling rapid and reversible tuning of gene-regulatory circuits without the need for transcriptional changes [36]. However, under conditions of persistent oxidative stress, such as in smokers’ lungs, this redox-sensitive architecture may become detrimental: chronic G oxidation distorts TL structure and impairs the maturation of TS G-rich TL miRNAs, such as miR-139, thereby facilitating oncogenic signaling and carcinogenesis (Figure 4). This model may represent an evolutionary trade-off, providing regulatory plasticity under normal conditions while predisposing to pathological miRNA loss in chronically oxidizing environments.

**Figure 4:**
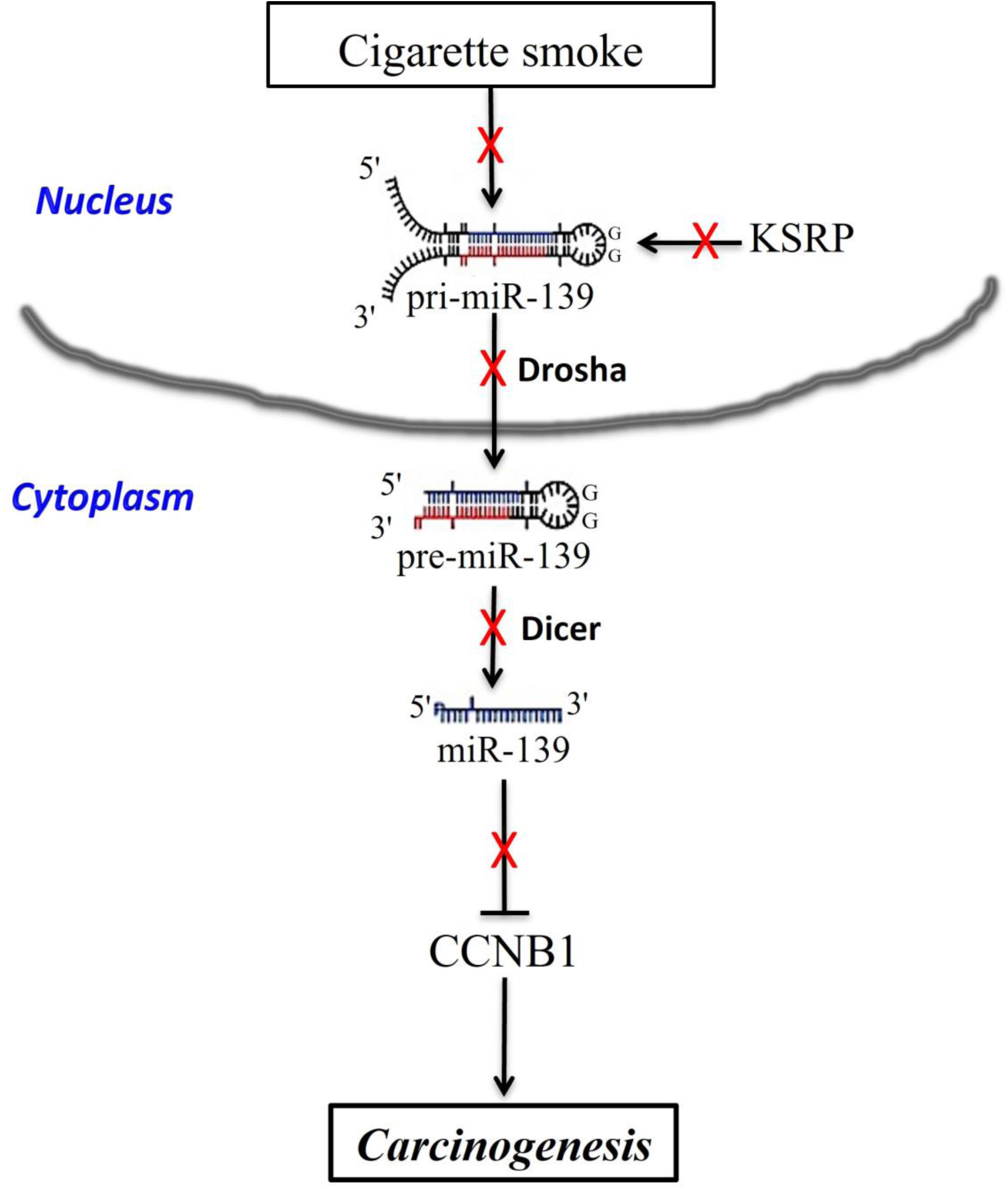
A flowchart illustrating the miRNA biogenesis pathway and its potential modulation by G content in TLs. Disruption of the processing of the TS miR-139 (indicated by red crosses) by carcinogens, such as cigarette smoke, leads to increased expression of its oncogenic target CCNB1 and contributes to carcinogenesis. The red arrow denotes upregulation, and the blue arrow denotes downregulation.

Several RBPs that are central to miRNA biogenesis, including Lin28 and KSRP, recognize G-rich motifs in TLs (GGAG and AGGGU, respectively) [37, 38]. These motifs serve as critical docking sites that facilitate high-affinity binding of RBPs to specific primary miRNA transcripts (pri-miRNAs), modulating microprocessor-dependent Drosha cleavage [39]. Structural studies have demonstrated that the exposure and flexibility of G-rich loops contribute to selective RBP recognition, allowing for context-dependent regulation of miRNA maturation [40]. Importantly, these G-rich motifs are particularly vulnerable to chemical modifications. Oxidative lesions such as 8-oxoG, as well as covalent adduct formation with reactive metabolites or pollutants, can alter the local conformation or base-pairing capacity of TLs [41, 42]. Such changes may reduce the affinity of RBPs for their cognate motifs, leading to inefficient processing of TS miRNAs and contributing to oncogenic signaling (Figure 4). Thus, the structural integrity of G-rich TL motifs is not only essential for normal miRNA biogenesis but may also represent a regulatory node susceptible to environmental or pathological redox stress.

Our findings provide an example of how the suppression of TS G-rich TL miR-139 may fail to counteract oncogenic drivers such as CCNB1 and highlight the therapeutic potential of stabilizing or restoring structurally vulnerable TS miRNAs, as recently demonstrated for the lung cancer-related miR-3196 [43]. Our study thus offers a conceptual framework for leveraging TL structural features in future therapeutic development, for example, by using synthetic miRNA mimics engineered with TL modifications that preserve regulatory function, reduce oxidative susceptibility, and potentially restore TS miRNA expression in lung cancer.

## CONCLUSIONS

Together with earlier observations, these results position TL G enrichment as a meaningful structural signature of TS miRNAs in lung cancer. By integrating miRNome-wide structural scoring with large-scale literature curation and functional annotation, this work provides evidence that TL G composition is not merely a passive sequence feature but a determinant of miRNA stability and anti-oncogenic function in the lung. Future studies should experimentally dissect the biochemical consequences of G oxidation in TLs and explore structural and chemical strategies to protect vulnerable TS miRNAs as a path toward miRNA-based therapeutics in lung cancer.

## Acknowledgements

None

## Funding

The authors declare that no funds, grants, or other support were received during the preparation of this manuscript.

## Supplementary Material

The following supplementary data are available for this study online:

**Supplementary Table 1:** A list of 955 miRNAs with pre-miRNA TL sequences (retrieved from the miRBase database), G and GG percents and G enrichment empirical p-values.

**Supplementary Table 2:** A list of 42 G-rich TL miRNAs and 17 G-free TL miRNAs, their annotation confidence (according to miRBase database), and the PubMed Identifier (PMID) numbers of PubMed Database references from the years 2015-2025 describing their function as oncomiRs (O) or tumor suppressor (TS) miRNAs. N.D = No Data.

**Supplementary Table 3:** A list of 61 experimentally validated target genes of the 42 TL G-rich TL miRNAs in lung cancer subtypes. Shown are the PubMed Identifier (PMID) numbers of PubMed Database references from the years 2015-2025 describing their function as oncomiRs (O) or tumor suppressor (TS) miRNAs.

## Author contributions

Conceptualization, A.C., M.A.B-A., Y.S.; validation, A.C., and Y.S.; investigation, A.C.; data curation, A.C. and Y.S.; writing—original draft preparation, A.C., and M.A.B-A.; writing—review and editing, A.C., and M.A.B-A.; visualization, A.C., M.A.B-A., and Y.S.; supervision, A.C., M.A.B-A., and Y.S. All authors have read and agreed to the published version of the manuscript.

## Declaration

### Competing interests

The authors declare no competing interests.

### Consent to Participate and Publication

There are no conflicts of interest among the authors regarding authorship or publication of this article. We confirm that this manuscript is original, has not been published previously, and is not under consideration elsewhere.

### Ethical Approval

The authors declare that they have followed the ethical guidelines outlined by the journal and that this study was conducted with utmost respect for the rights and dignity of all participants. However, there is no requirement for formal ethical approval, as results are based on the in-silico studies with no human involvement.

## Notes

### Competing Interest Statement

The authors have declared no competing interest.

